# NRF2 activates a partial Epithelial-Mesenchymal Transition and is maximally present in a hybrid Epithelial/Mesenchymal phenotype

**DOI:** 10.1101/390237

**Authors:** Federico Bocci, Satyendra C Tripathi, Samuel A Vilchez Mercedes, Jason T George, Julian P Casabar, Pak Kin Wong, Samir M Hanash, Herbert Levine, José N Onuchic, Mohit Kumar Jolly

## Abstract

The Epithelial-Mesenchymal Transition (EMT) is a key process implicated in cancer metastasis and therapy resistance. Recent studies have emphasized that cells can undergo partial EMT to attain a hybrid epithelial/mesenchymal (E/M) phenotype - a cornerstone of tumour aggressiveness and poor prognosis. These cells can have enhanced tumour-initiation potential as compared to purely epithelial or mesenchymal ones and can integrate the properties of cell-cell adhesion and motility that facilitates collective cell migration leading to clusters of Circulating Tumour Cells (CTCs) - the prevalent mode of metastasis. Thus, identifying the molecular players that can enable cells to maintain a hybrid E/M phenotype is crucial to curb the metastatic load. Here, using an integrated computational-experimental approach, we show that the transcription factor NRF2 can prevent a complete EMT and instead stabilize a hybrid E/M phenotype. Knockdown of NRF2 in hybrid E/M non-small cell lung cancer cells H1975 and bladder cancer cells RT4 destabilised a hybrid E/M phenotype and compromised the ability to collectively migrate to close a wound in vitro. Notably, while NRF2 knockout simultaneously downregulated E-cadherin and ZEB-1, overexpression of NRF2 enriched for a hybrid E/M phenotype by simultaneously upregulating both E-cadherin and ZEB-1 in individual RT4 cells. Further, we predict that NRF2 is maximally expressed in hybrid E/M phenotype(s) and demonstrate that this biphasic dynamic arises from the interconnections among NRF2 and the EMT regulatory circuit. Finally, clinical records from multiple datasets suggest a correlation between a hybrid E/M phenotype, high levels of NRF2 and its targets and poor survival, further strengthening the emerging notion that hybrid E/M phenotype(s) may occupy the ‘metastatic sweet spot’.

## Introduction

Epithelial-Mesenchymal Transition (EMT) is a hallmark of physiological and pathological processes such as embryonic development, wound healing and tumour progression [1]. During EMT, cells typically lose cell-cell adhesion and gain the traits of migration and invasion. However, EMT is not a binary process; instead, cells can also attain one or more hybrid epithelial/mesenchymal (E/M) phenotypes that may be more stem-like, metastatic and aggressive than those in completely epithelial or mesenchymal ones [1, 2]. Cells in hybrid E/M phenotype(s) display a mix of epithelial and mesenchymal traits, thereby enabling collective cell migration leading to the formation of clusters of Circulating Tumour Cells (CTCs) that can form a large majority of metastases, and correlate with poor clinical outcome [3, 4].

Mechanisms enabling cancer cells to maintain hybrid E/M phenotype(s), however, remain largely unclear. Recent experimental and computational studies have suggested that a set of ‘phenotypic stability factors’ (PSFs) such as GRHL2, OVOL2, ΔNp63α and NUMB can help cells in maintaining a hybrid E/M phenotype by preventing ‘cells that have gained partial plasticity’ from undergoing a full EMT [5–10] and thus enabling collective cell migration. Knockdown of these PSFs drove the cells to a complete EMT *in vitro* and *in vivo,* while their exogenous overexpression can drive Mesenchymal-Epithelial Transition (MET) – the reverse of EMT [11]. Thus, the relative levels of PSFs and canonical EMT-inducing transcription factors (EMT-TFs) modulate the epithelial-mesenchymal plasticity of cells. A recent study highlighted the role of the transcription factor nuclear factor E2-related factor 2 (NFE2L2) – commonly referred to as NRF2 – in preventing cells from undergoing a full EMT during injury-induced wound healing [12]. Hence, we hypothesized that NRF2 can also act as a PSF.

NRF2 connects with EMT circuitry through the following links: a) NRF2 can inhibit EMT-TF Snail [13], b) members of miR-200 family – gatekeepers of an epithelial phenotype [14] – can increase NRF2 levels by targeting KEAP1 for ubiquitination and hence preventing the cytoplasmic degradation of NRF2 [15], and c) nuclear accumulation and transcriptional activity of NRF2 can be restricted by E-cadherin [16] (Fig. 1A); thus, it is possible that ZEB1 – a key EMT-TF that represses E-cadherin directly [17] – can effectively increase the nuclear levels and activity of NRF2.

**Figure 1.**
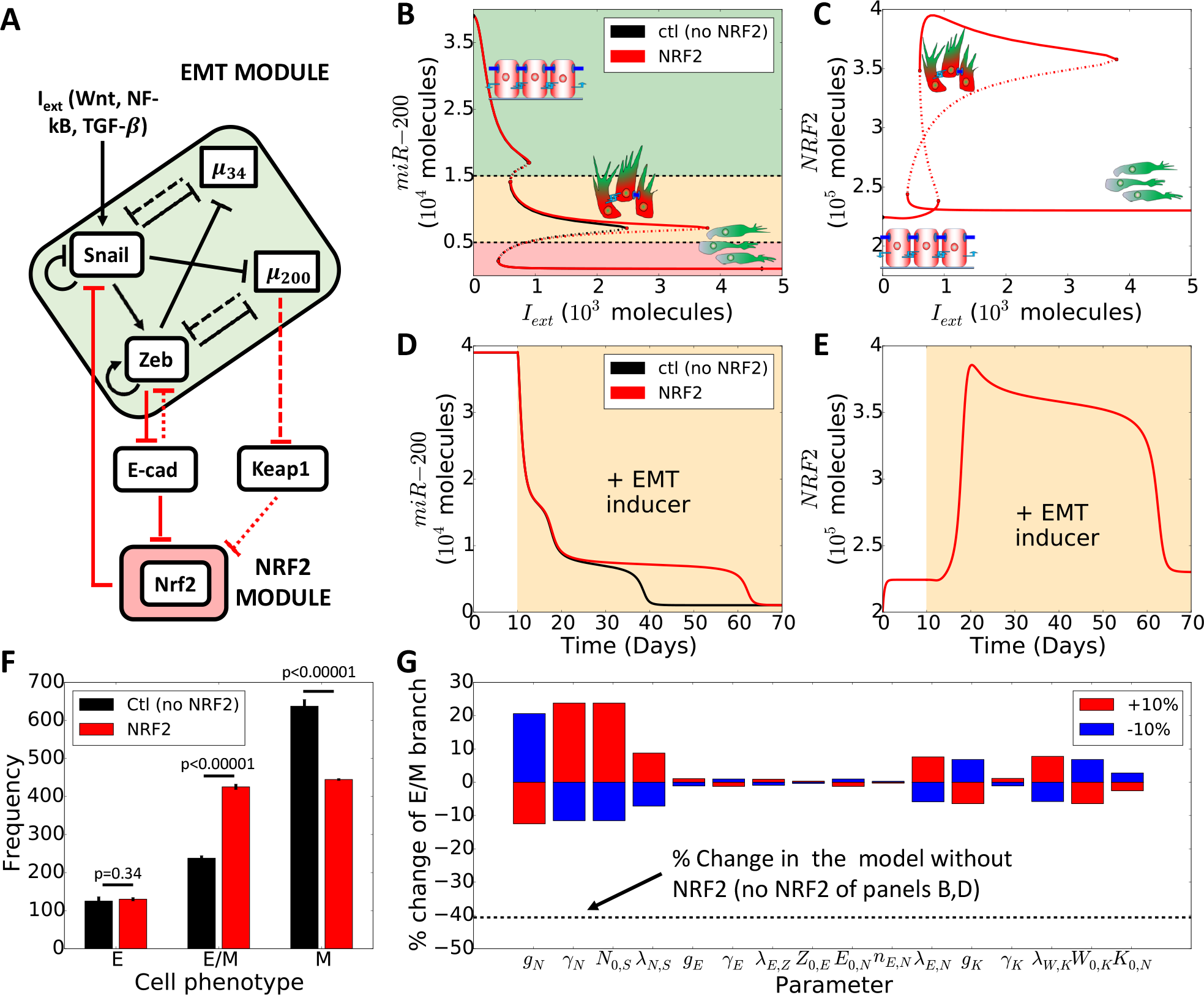
NRF2 stabilizes a hybrid E/M phenotype in presence of an external EMT-inducing signal. **(A)** Schematic of EMT-NRF2 intracellular coupled regulatory network. Red arrows represent NRF2-related interactions that are not considered in the control case (i.e. NRF2 absent). Solid arrows represent transcriptional activation and solid bars stand for transcriptional inhibition. Dotted lines depict micro-RNA mediated translational inhibition. KEAP1 inhibits NRF2 by facilitating degradation while E-cadherin inhibits ZEB via β-catenin (fine-dotted arrows). **(B)** Bifurcation diagram of miR-200 levels as a function of I_ext_. Curves denote the value of miR-200 upon equilibration, where continuous lines stand for stable steady states and dotted lines represent unstable solutions (black for the control case, red for NRF2-dependent case). Cartoons and colour shadings depict epithelial (green), hybrid E/M (yellow) and mesenchymal (red) phenotypes. **(C)** Bifurcation diagram of NRF2 levels as a function of I_ext_. **(D)** Temporal dynamics of miR-200 levels in a cell starting in an epithelial phenotype, when exposed to a high level of I_ext_ = 5000 molecules (orange-shaded region). **(E)** Temporal dynamics of NRF2 levels, for case shown in D). **(F)** Number of independent simulations of a cell starting from random initial conditions and exposed to varied level of I_ext_ chosen randomly in [0, 5000] molecules that resulted in an E, hybrid E/M, or M phenotype, respectively. Error bars indicate the standard deviation of three independent simulation sets (n=1000 each) for the control case (black) and the case with NRF2 (red). p-values were computed using a two-tailed paired t-test. **(G)** Variation in the interval of stability of the hybrid E/M phenotype (see panel 1B) in response to a 10% variation of the model parameters. Black dotted line indicates the relative change in the stability of hybrid E/M phenotype for the control case (NRF2 absent).

Elucidating the emergent dynamics of the crosstalk among EMT and NRF2 signalling pathways requires constructing a mechanism-based quantitative model. Here, we incorporate the abovementioned connections of NRF2 into our previously developed mathematical framework for EMT signalling [18]. Our model predicts that NRF2 can act as a PSF for a hybrid E/M phenotype. To validate this prediction, we knocked down NRF2 in H1975 non-small cell lung cancer cells (NSCLC) [6] and RT4 bladder cancer cells that display a hybrid E/M phenotype stably over multiple passages *in vitro.* Knockdown of NRF2 drove the cells towards a mesenchymal phenotype at a molecular and morphological level, hence showing the role of NRF2 in maintaining a hybrid E/M phenotype. Conversely, overexpressing NRF2 through DL-Sulforaphane (SFN) in RT4 bladder cancer cells resulted in an increase of both E-cadherin and ZEB1, hence confirming the association between NRF2 and a hybrid E/M phenotype. Furthermore, we suggest that NRF2 exhibits a biphasic response during EMT progression and reaches its maximal levels when the cell is in a hybrid E/M phenotype. This biphasic dynamics emerges from the topology of the network formed between NRF2 and the core EMT circuit. Finally, we found correlations among elevated expression levels of NRF2 and its transcriptional targets, a hybrid E/M phenotype and poor patient survival in multiple clinical datasets, hence strengthening the emerging notion that a hybrid E/M phenotype, but not necessarily a completely mesenchymal one, can aggravate tumour progression.

## Results

### Mathematical modelling predicts that high NRF2 levels can stabilize and are a hallmark of a hybrid E/M phenotype

To elucidate the role of NRF2 in mediating EMT, we first investigated the crosstalk between NRF2 and a core network that regulates EMT (Fig. 1A). This core EMT network includes two families of EMT-TFs, SNAIL and ZEB, and two families of microRNAs that inhibit EMT, miR-200 and miR-34 [18] (Fig. 1A). The intracellular dynamics of NRF2-EMT circuit was described via a system of coupled ordinary differential equations (ODEs) (see Methods) to investigate how NRF2 affects the response of a cell when it is exposed to an external EMT-inducing signal.

First, we calculated the bifurcation diagram of cellular EMT phenotypes in response to different levels of an external EMT-inducing signal (*I*_*ext*_). For increasing values of *I*_*ext*_, cells switch from an epithelial phenotype to a hybrid E/M phenotype to a mesenchymal phenotype, as displayed in the decreasing levels of miR-200 (Fig. 1B) and increasing levels of EMT-TFs (Supplementary Fig. S1A). In the presence of NRF2, a cell requires a much stronger stimulus to undergo a complete EMT and instead maintains a hybrid E/M phenotype over a much broader range of the external signal *I*_*ext*_, as compared to the control case – i.e. NRF2 absent (Fig. 1B, compare the red and black curves). This result is quite robust upon variation of the coupling strength among NRF2 and the EMT circuit (Supplementary Fig. S1B-C). Intriguingly, the model predicts that cells in a hybrid E/M phenotype have higher levels of NRF2 as compared to those in epithelial or mesenchymal ones (Fig. 1C). Such behaviour emerges because both miR-200, an EMT-inhibiting microRNA, and ZEB, an EMT-TF, inhibit molecular players that, in turn, inhibit NRF2 (KEAP1 and E-cadherin, respectively). Therefore, NRF2 receives an indirect activation from both mir-200 and ZEB, thereby leading to higher levels in a hybrid E/M phenotype.

Next, we investigated the temporal dynamics of a cell in response to the EMT-inducing signal (*I*_*ext*_). In absence of NRF2, a cell starting from an epithelial state (i.e. high miR-200, low ZEB) first undergoes a partial EMT (i.e. transition to a hybrid E/M phenotype) and later attains a mesenchymal phenotype (Fig 1D, black curve). The presence of NRF2, however, delays the transition to the fully mesenchymal state and maintains the hybrid E/M phenotype much longer (Fig. 1D, red curve). Concurrently, NRF2 increases during partial EMT and reduces once the cell attains a complete mesenchymal phenotype (Fig. 1E).

Our model therefore suggests that NRF2 prevents the transition from a hybrid E/M phenotype to a M phenotype, but not the transition from an E phenotype to a hybrid E/M phenotype. This differs from the behaviour of other phenotypic stability factors (PSFs), such as Numb, that inhibits EMT independently from the cell’s specific phenotype (Supplementary Fig. 2). To further investigate this hypothesis, we performed multiple simulations exposing the cell to random levels of EMT-inducing signal *I*_*ext*_ and counted the frequency of attaining E, E/M and M phenotypes, upon full equilibration of the system. We found an increased fraction of hybrid E/M cells and decreased fraction of M cells in presence of NRF2, while the frequency of the E phenotype remains unaltered (Fig. 1F), further strengthening our hypothesis.

Finally, to identify the robustness of the observed behaviour, we performed a sensitivity analysis where we varied the parameters of the model by ± 10% – one parameter at a time – and measured the corresponding change in the range of values of *I*_*ext*_ enabling a stable hybrid E/M phenotype (i.e. interval of x-axis where the hybrid E/M state is stable in Fig. 1B, as shown by the red solid middle branch). For most parameters, the relative change in this range of values of *I*_*ext*_ is small, indicating that the model is largely robust. On the other hand, the parameters related to NRF2 production, degradation, and interaction with SNAIL showed a stronger sensitivity (Fig. 1G). These sensible parameters control the overall effect of NRF2 on EMT, and thus affect the stability of the hybrid E/M phenotype more directly. This decrease, however, is much weaker as compared to the control case (dotted line on Fig. 1G, i.e. NRF2 absent). Overall, these results suggest that NRF2 can act as a ‘phenotypic stability factor’ (PSF) for a hybrid E/M phenotype and its enhanced levels can be considered as a hallmark of the hybrid E/M phenotype.

### NRF2 knockdown in H1975 NSCLC cells destabilizes a hybrid E/M phenotype *in vitro*

To investigate our prediction that NRF2 can act as a PSF for a hybrid E/M phenotype, we knocked down NRF2 using siRNAs in non-small cell lung cancer (NSCLC) H1975 cells that display a hybrid E/M phenotype stably *in vitro* over multiple passages [6].

We observed that knockdown of NRF2 increases the EMT-TFs ZEB1 and SNAIL and mesenchymal marker vimentin at both mRNA and protein levels, and decreased the mRNA levels of E-cadherin (in particular siNRF2 #2). The protein levels of E-cadherin, however, did not change significantly, potentially due to its long half-life [19] (Fig. 2A,B). These observations suggest that NRF2 knockdown (NRF2-KD) can drive cells from a hybrid E/M phenotype to a mesenchymal phenotype, as predicted by the mathematical modelling (Fig. 2C). Consistently, bright-field microscopy revealed a cellular morphology characterized by more spindle-shaped traits characteristic of mesenchymal cells upon knockdown of NRF2 (Fig. 2D). Further, H1975 cells partially lose the ability to migrate after NRF2 knockdown in transwell migration assays (Fig. 2E and 2G, left). Similarly, NRF2 knockdown compromised the cells’ ability to move collectively and close the wound. While control cells almost exclusively moved collectively while maintaining physical cell-cell contacts, KD-NRF2 compromised wound closure and gave rise to a subpopulation of cells that migrated solitarily (Fig. 2F and 2G, right), thus validating our hypothesis that a hybrid E/M cellular phenotype is being destabilized and cells progress towards a completely mesenchymal state.

**Figure 2.**
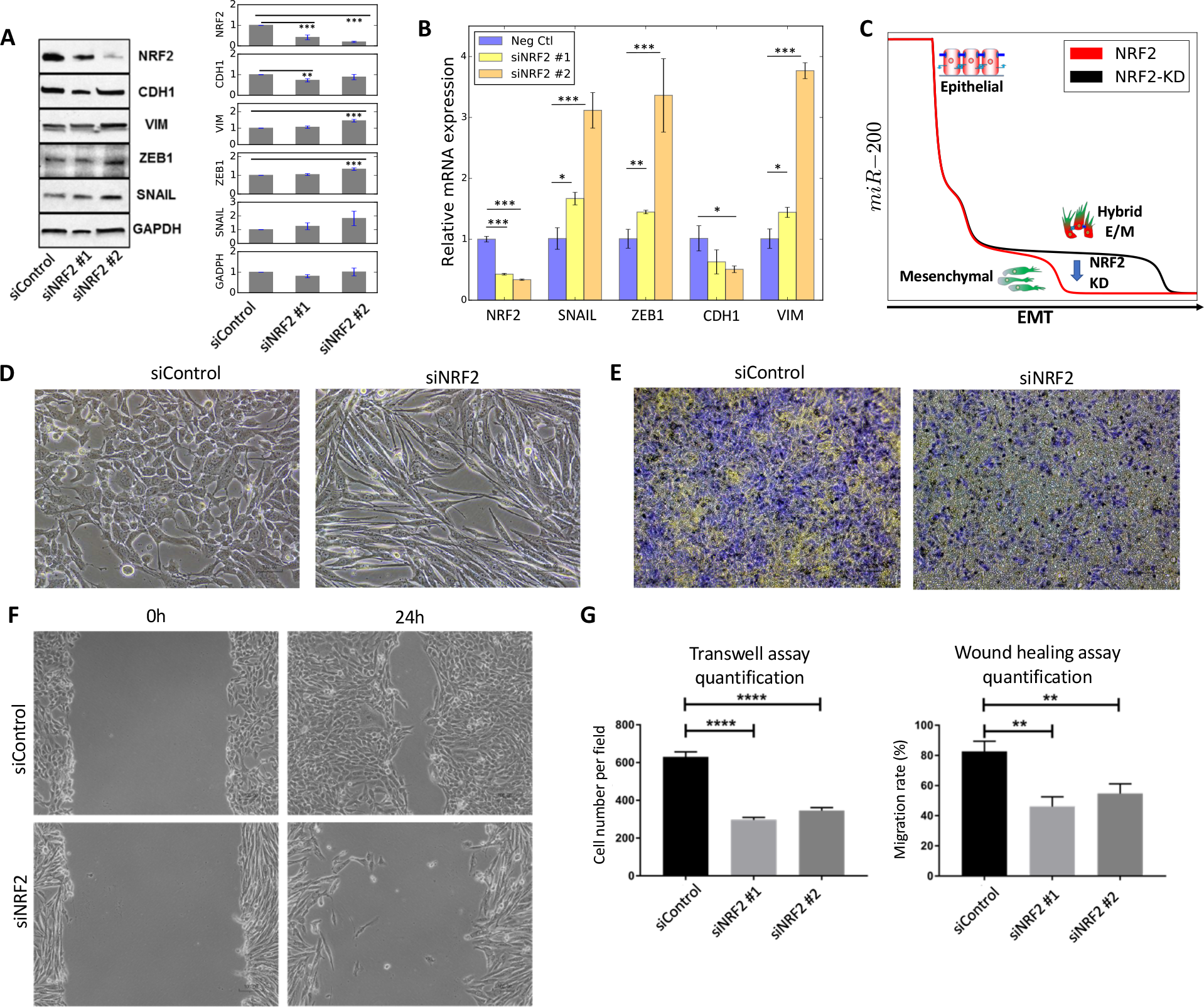
Knockdown of NRF2 induces a progression towards a complete EMT in H1975 cells. **(A)** Western blot for NRF2, CDH1, VIM, ZEB1, SNAIL in H1975 cells. GAPDH is used as a loading control. Left column represents the control siRNA, second and third columns represent two independent siRNAs against NRF2. Bar chart shows the average over 3 biological replicates. **(B)** mRNA levels of NRF2, SNAlL, ZEB, CDH1 (E-cadherin) and VIM (vimentin) for siRNA control (Neg Ctl) and cells treated with siRNA against NRF2. **(C)** Temporal profile of miR-200 in response to an EMT-inducing signal in presence of NRF2 (red, same as ‘NRF2’ case Fig. 1D) and upon NRF2-KD (black) as predicted by mathematical modelling. NRF2 knockdown is modelled as a 90% reduction in the production rate constant of NRF2, and compromises the temporal stability of the hybrid E/M phenotype as depicted by the blue arrow. **(D)** Bright-field morphology images for control (left) and NRF2-knockdown (right) H1975 cells. **(E)** Transwell migration assay showing trends of collective migration (left, Neg ctl) and individual migration (right, siNRF2). Violet colour represents cells while yellow colour represents membrane **(F)** Wound healing/scratch assay for control (neg ctl, top) and NRF2-KD (siNRF2, bottom) H1975 cells (magnification 100x). **(G)** Quantification of transwell migration assay (left) and wound healing assay (right). Wound healing assay was quantified in terms of the ability to close the wound. *p < 0.05, **p<0.005, ***p<0.001 using two-tailed paired t-test. For (D-E-F), siNRF2 #2 was used for NRF2-KD.

Overall, NRF2 knockdown shows a progression to a complete EMT both at molecular and morphological levels, thus confirming that NRF2 can act as a PSF for a hybrid E/M phenotype.

### NRF2 knockout in RT4 bladder cancer cells destabilises a hybrid E/M phenotype *in vitro*

To gain further confidence in our hypothesis that NRF2 can stabilise a hybrid E/M phenotype, we obtained a Crispr/Cas9 NRF2 cell pool knockout (KO) in a different cell line - bladder cancer RT4 cells that express a stable hybrid E/M phenotype over multiple passages *in vitro* as shown by immunostaining of E-cadherin and ZEB-1 (Supplementary Fig. S3). The knockout of NRF2 was confirmed by downregulation of NRF2 and the NRF2 downstream gene target HO-1 (Fig. 3A-B). Strikingly, NRF2-KO decreases the expression of both the epithelial marker E-cadherin and the mesenchymal marker ZEB-1 (Fig. 3C-E, H), therefore suggesting that NRF2-KO destabilises the hybrid E/M phenotype. Moreover, similar to the case of H1975 NSCLC cells, NRF2-KO severely restricts the ability of RT4 cells to migrate and collectively close a wound (Fig. 3F).

**Figure 3.**
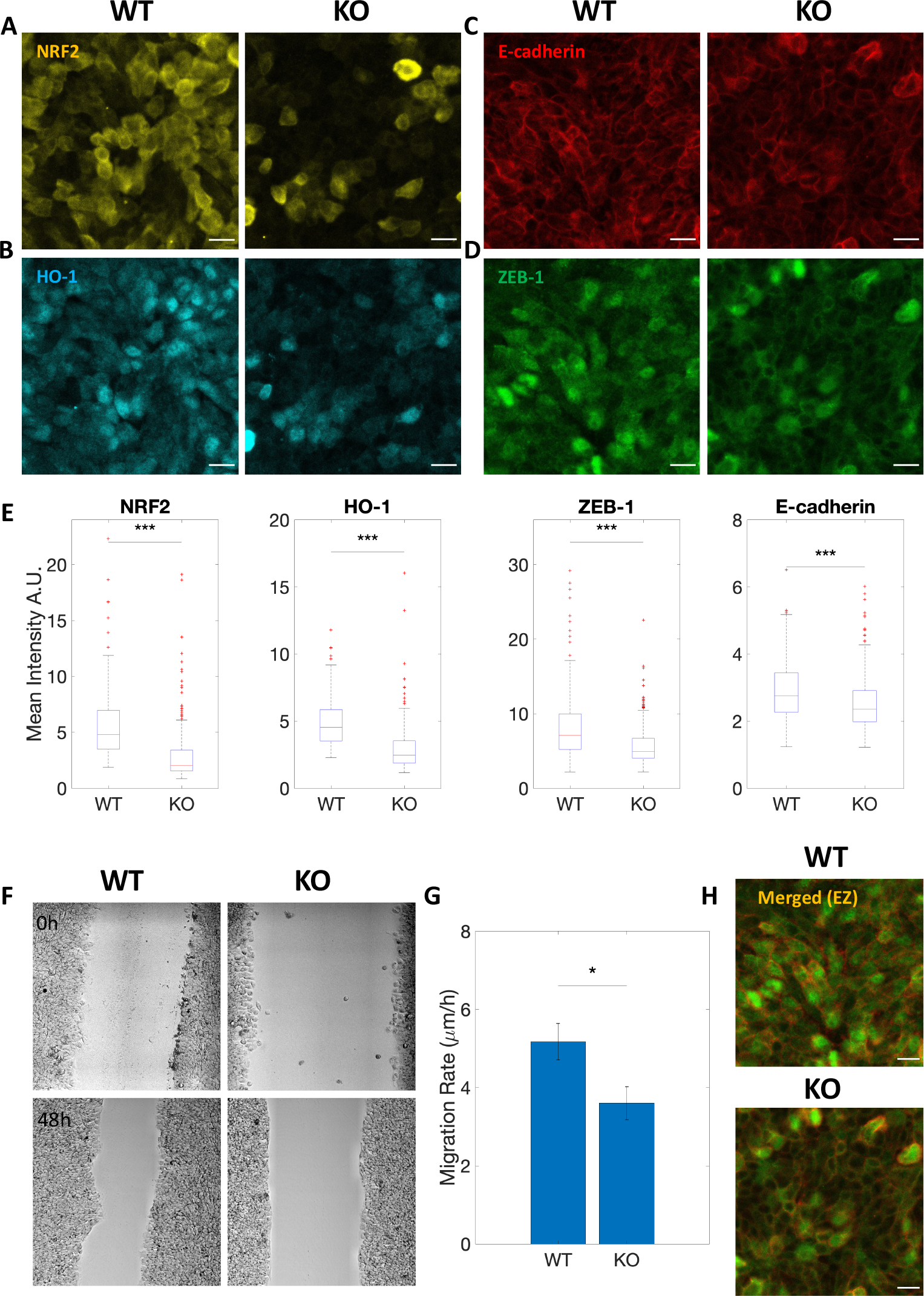
NRF2 knockout alters the hybrid E/M phenotype in RT4 bladder cancer cells in vitro. **(A-D)** NRF2, HO-1, E-cadherin, and ZEB-1 protein expression through immunocytochemistry for RT4 wild type (WT) and Crispr/Cas9 NRF2-KO (KO), respectively. Scale bar: 20 μm. **(E)** Box plot of WT vs KO for NRF2, HO-1, E-cadherin and ZEB-1. Data are shown as boxplot of cell intensity in arbitrary units (A.U.). The data was analyzed in MATLAB and the Mann-Whitney-U test was used to obtain the p-value, where ‘***’ represents a p-value < 0.001. At least 250 cells per condition were measured in each experiment. **(F)** Brightfield images of WT vs KO at 0 hours and 48 hours after wound healing (10x objective in Leica TCS SP8 microscope). **(G)** Bar graph of WT vs KO showing decreased migration rate for KO. Data is shown as mean with standard error mean (SEM). The data was analyzed in MATLAB and the Student’s two-tailed t-test was used to obtain the p-value, where ‘*’ represents a p-value <0.05. **(H)** Merged view of E-Cadherin and ZEB-1 for WT vs KO condition. All Images are representative of at least 3 independent experiments.

Overall, observations from H1975 NSCLC and RT4 bladder cancer cells indicate that NRF2 is required to stabilise a hybrid E/M phenotype, and that the loss of NRF2 can destabilise such hybrid E/M phenotype(s).

### SFN-induced NRF2 overexpression in RT4 cells enhances the hybrid E/M phenotype *in vitro*

To strengthen our hypothesis regarding the role of NRF2 in maintaining a hybrid E/M phenotype, we pharmacologically activated NRF2 in RT4 bladder cancer cells through SFN [13, 20]. We observed activation of NRF2 and HO-1 upon SFN treatment (7.5 μM, 24h) (Fig. 4A-B). Remarkably, upregulating NRF2 from its basal level via SFN resulted in a significant increase of both the epithelial marker E-cadherin and the mesenchymal marker ZEB-1 (Fig. 4C-F), i.e. the opposite outcome of NRF2-KO in RT4 cells. This observation is consistent with the prediction from the mathematical model. In fact, overexpressing NRF2 significantly increased the fraction of hybrid E/M cells, as compared to a case with normal levels of NRF2 (Fig. 4G). These observations lend further support to the proposed role of NRF2 in maintaining/stabilizing a hybrid E/M cell phenotype.

**Figure 4.**
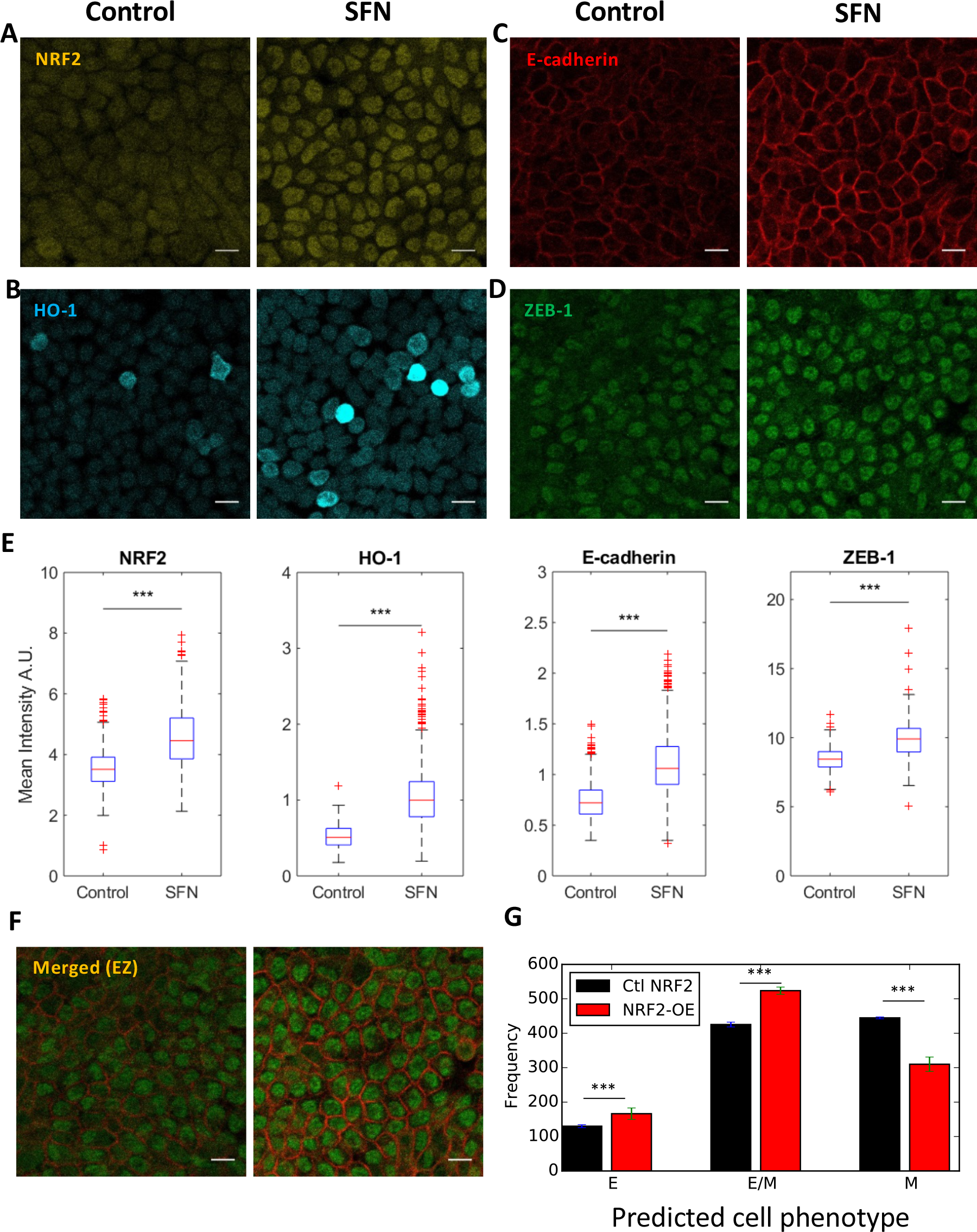
NRF2 upregulation enhances the hybrid E/M phenotype in vitro. **(A-D)** Control and SFN treatment for NRF2, HO-1, E-cadherin and ZEB-1. Scale bar: 20 μm. **(E)** Box plot of Control vs SFN for NRF2, HO-1, E-cadherin and ZEB-1. Data are shown as a boxplot of cell intensity in arbitrary units (A.U.). The data was analyzed in MATLAB and the Student’s two-tailed t-test was used; ‘***’ denotes a p-value < 0.001. At least 1000 cells per condition were measured in each experiment. Images are representative of at least 3 independent experiments. **(F)** Merged view of E-cadherin and ZEB-1. Scale bar: 20 μm. **(G)** Fraction of E, E/M, and M cells exposed to varied level of I_ext_ chosen randomly in [0, 5000] molecules for a control case with NRF2 (black) and 4-fold NRF2 overexpression (red). Cell classification was done upon full equilibration. Error bars indicate the standard deviation of three independent simulation sets (n=1000 independent cells for each set). p-values were computed using a two-tailed paired t-test.

### Enhanced levels of NRF2 in a hybrid E/M phenotype emerge from the underlying network topology

To understand the detailed origin of the connection between NRF2 and EMT, we applied a recently developed computational analysis method - Random Circuit Perturbation (RACIPE) [21]. RACIPE takes the topology of a regulatory network as its input, and generates an ensemble of mathematical models in which parameters are randomly chosen. For every set of randomly chosen parameters, the algorithm returns the levels of all the considered variables upon full equilibration. Thus, it identifies robust gene expression profiles that can be expected from a given network topology. For the circuit we considered, the RACIPE output is an array containing the levels of miR-34, miR-200, ZEB, SNAIL, E-Cadherin, KEAP1, and NRF2. These levels are presented by the algorithm using a logarithmic scale and can therefore assume negative values.

After evaluating the NRF2-EMT circuit for 10000 randomly chosen sets of parameters, we performed a clustering of the obtained solutions via a K-means algorithm for K=3. Importantly, NRF2 was not considered for the clustering to allow an unbiased evaluation of the association of NRF2 levels with various clusters. Among the three clusters, one cluster (or group of solutions) is characterized by high expression of microRNAs and E-cadherin and low expression of EMT-TFs, and could thus be associated to an epithelial phenotype (Fig. 5A, bottom). The second cluster exhibits high expression of EMT-TFs and low microRNAs – signature for a mesenchymal phenotype (Fig. 5A, top). Finally, the third group has intermediate to high levels of both microRNAs and EMT-TFs and is thus interpreted as a hybrid E/M phenotype (Fig. 5A, center). Importantly, when plotted on miR-200 and ZEB axes, these clusters segregate elegantly and occupy different regions, suggesting that miR-200 and ZEB dynamics may play crucial roles in driving the emergence of these three phenotypic clusters (Fig. 5B) [18, 22].

**Figure 5.**
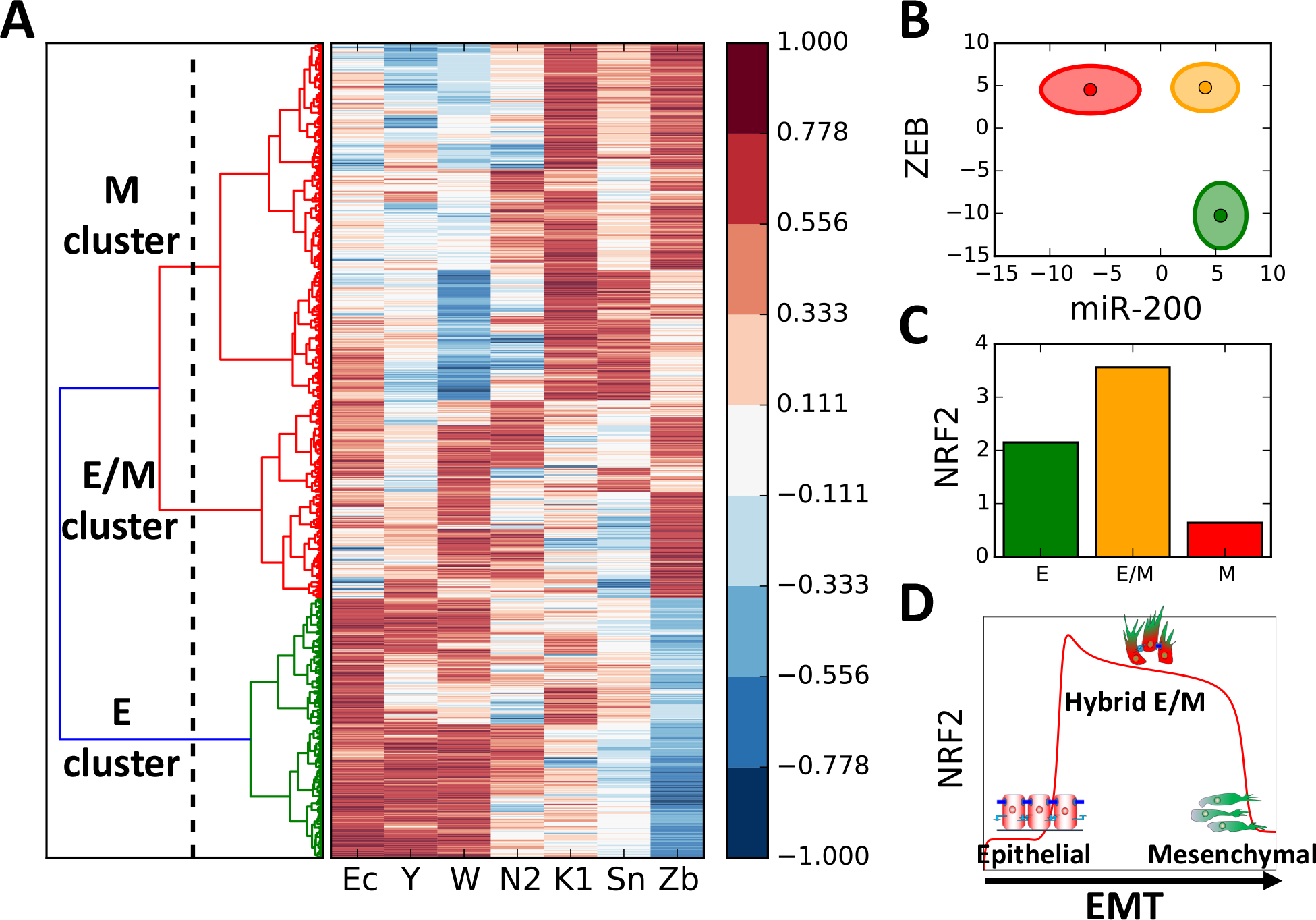
Random Circuit Perturbation of the EMT-NRF2 circuit. **(A)** Right: gene expression profile of the states predicted by RACIPE. Each row represents a solution obtained with a different randomly chosen combination of model’s parameters presented here as an array of the model’s variables (Ec: E-cadherin, Y: miR-34, W: miR-200, N2: NRF2, K1: Keap1, Sn: SNAIL, Zb: ZEB). All gene expression levels are normalized between −1 and 1 to help eye comparison. Left: clustering of the solutions and threshold for K-means clustering (vertical dotted line leaving n=3 groups of solutions). **(B)** Representation of the three clusters in the miR-200/ZEB plane. The centres of the ellipses represent the cluster average while the axes quantify the cluster’s standard deviation. Solutions belonging to the epithelial, hybrid E/M and mesenchymal clusters are plotted in green, yellow, and red, respectively. **(C)** Average level of NRF2 in the epithelial, hybrid E/M and mesenchymal clusters. **(D)** Temporal profile of NRF2 in response to an EMT-inducing signal as predicted by mathematical modelling (Fig. 1E) with relevant cartoons added.

Next, we computed average levels of NRF2 in these clusters, and showed that the hybrid E/M cluster expresses higher levels of NRF2 (Fig. 5C and Supplementary Fig. S4A) as compared to epithelial or mesenchymal clusters, hence supporting our model’s prediction (Fig. 5D). This analysis highlights that the maximal level of NRF2 is not an artefact enabled by a specific set of parameters, but instead emerges from the connections, or architecture of the network of interactions among NRF2 and the EMT module.

To better understand the underlying topological features enabling maximal levels of NRF2 in hybrid E/M state, we performed *in silico* simulations where either the inhibition of NRF2 by KEAP1 (KEAP1-NRF2 KD) or that by E-cadherin (Ecad-NRF2 KD) was deleted. KEAP1-NRF2 KD maintained high NRF2 levels in the hybrid E/M cluster, but Ecad-NRF2 KD resulted in higher NRF2 in the epithelial cluster (Supplementary Fig. S4B-C). Consistently, reducing the effect of KEAP1-mediated inhibition of NRF2 in our EMT-NRF2 mathematical model (shown in Fig 1A) maintained high NRF2 in the hybrid E/M state, while reducing E-Cadherin-mediated inhibition of NRF2 reduced the difference in NRF2 levels between the E and hybrid E/M phenotypes (Supplementary Fig. S5). This asymmetry can be perhaps attributed to a larger number of connections among ZEB, SNAIL, E-cadherin and NRF2.

### Clinical data suggests correlation between worse survival, high NRF2 levels and a hybrid E/M phenotype

The association of NRF2 with hybrid E/M phenotype(s) suggests addressing the role of NRF2 in a clinical context. Recent studies have emphasized that a hybrid E/M phenotype can be considered as a hallmark of tumour aggressiveness [1, 2] due to multiple reasons – the enabling of CTC dissemination as clusters instead of individual cells (typically mesenchymal), and the enhancement of tumour-initiation potential. Consistently, several previously discovered PSFs correlate with a poor clinical outcome. Among other examples, GRHL2 was associated with poor survival in different breast cancer subtypes [23] and Numb/Numbl correlated with a worse clinical outcome in lung and ovarian cancer [8].

We analysed NRF2 expression levels in patients from multiple independent breast cancer datasets. Previous studies have highlighted association between worse survival and a hybrid E/M phenotype in breast cancer [6, 10]. We observed that NRF2 correlates with a worse clinical outcome both in terms of overall survival (Fig. 6A) and relapse-free survival (Fig. 6B-D), when these samples are divided by median. We further calculated the ‘EMT score’ for all these patients’ gene expression profiles using an inferential parameter developed by George *et al*. [24] based on the expression level of several key EMT regulators. These scores are calculated on a scale of 0 (highly epithelial) to 2 (highly mesenchymal), with the following categorization: epithelial (0<score<0.5), hybrid E/M (0.5<score<1.5) or mesenchymal (1.5<score<2). An EMT score distribution was thus reconstructed within each dataset for the ‘low NRF2’ group (NRF2<median) and the ‘high NRF2’ group (NRF2>median) (Fig. S5). From these EMT score probability distributions, it is possible to calculate the probability to observe an epithelial, hybrid E/M, or mesenchymal EMT score for patients belonging to the ‘high NRF2’ or the ‘low NRF2’ groups (see Supplementary Fig. S6). This analysis revealed that patient samples belonging to the ‘high NRF2’ group have a higher probability to display a hybrid E/M phenotype (0.5< EMT score <1.5), as compared to exhibiting either an E or M phenotype (Fig. 6E). Furthermore, patient samples characterized by a hybrid E/M phenotype (0.5< EMT score <1.5) have a higher probability of belonging to the ‘high NRF2’ category as compared to belonging to the ‘low NRF2’ one (Fig. 6E). Both these observations are consistent across the three datasets where we observed a significant association between high NRF2 and poor patient survival.

**Fig. 6.**
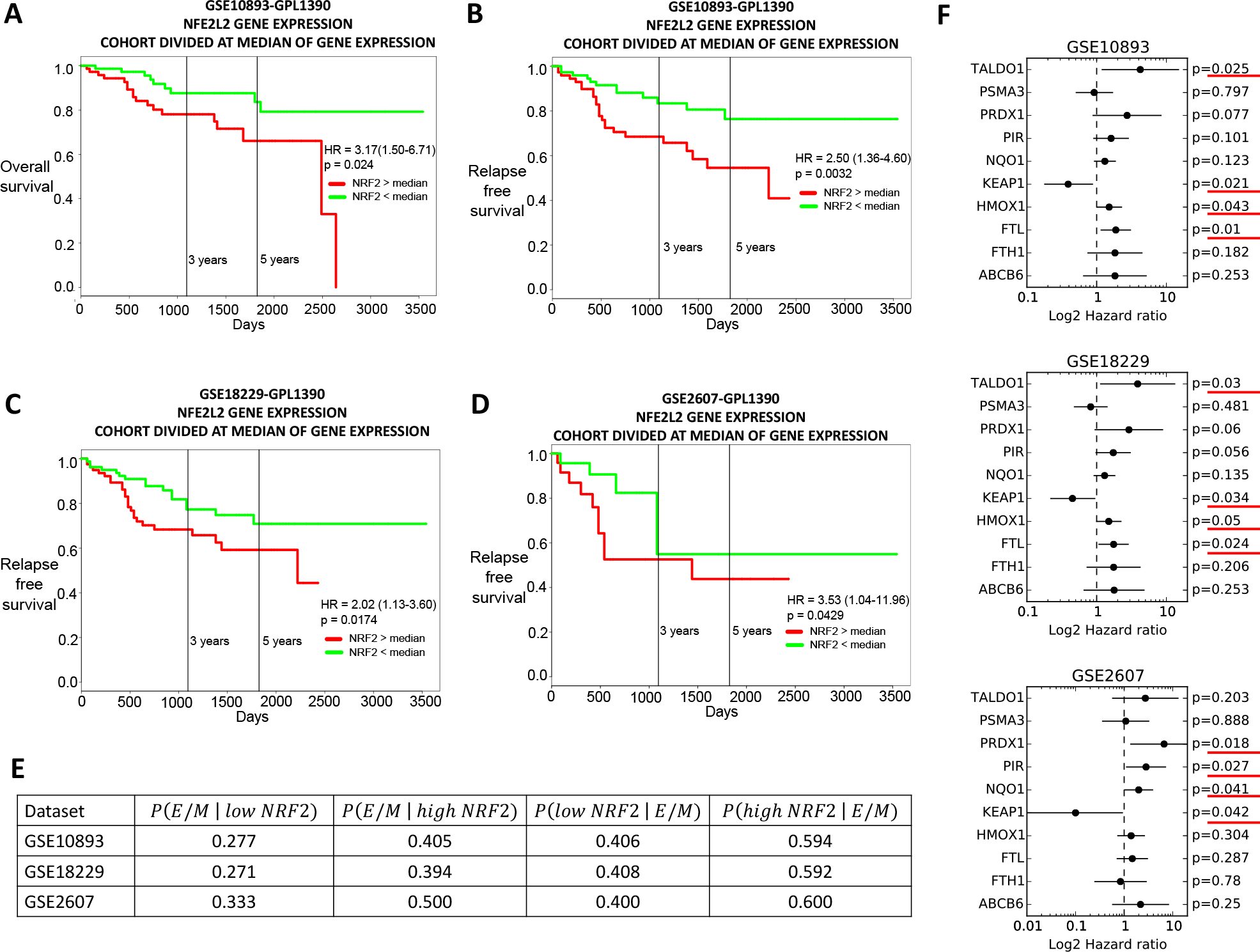
NRF2 correlates with a worse clinical outcome in cancer patients. **(A)** Overall survival for GSE10893. The group of patients with low NRF2 (green curve) performed better than the group with high NRF2 (red curve). **(B)** Relapse-free survival for GSE10893. **(C)** Relapse-free survival for GSE18229. **(D)** Relapse-free survival for GSE2607. **(E)** Left: conditional probability of exhibiting a hybrid E/M EMT score given a low NRF2 expression (second column) or a high NRF2 expression (third column). Right: conditional probability of a low NRF2 level (forth column) or a high NRF2 level (fifth column) given a hybrid E/M EMT score. **(F)** Hazard ratio, 95% confidence interval (error bars) and p value for several NRF2 downstream target genes for GSE10893 (top), GSE18229 (centre) and GSE207 (bottom).

Interestingly, NRF2 does not correlate with other phenotypic stability factors in the three considered datasets (Supplementary Fig. S6). This result can be explained by considering that previously identified ‘conventional’ PSFs, such as OVOL, GRHL2, or NUMB unilaterally prevent EMT and were maximally expressed in epithelial phenotype [6, 8, 25], while NRF2 is predicted to more specifically stabilise a hybrid E/M phenotype and being maximally present in the same.

Finally, due to the prominent regulation of NRF2 at post-transcriptional level by KEAP1 [15], we investigated the expression levels of NRF2 downstream targets [26]. High levels of many NRF2 downstream targets correlate with worse survival (hazard ratio > 1) in a statistically significant manner in the three considered datasets (Fig. 6F). Among those targets, only KEAP1 shows an opposite trend, i.e. its higher levels associated with better survival (Fig. 6F). This apparently counterintuitive finding can be rationalized because KEAP1 also acts as an inhibitor of NRF2, as also included in our mathematical model. Thus, put together, these results suggest a strong correlation among a hybrid E/M phenotype, high expression levels of NRF2 and its downstream targets, and a worse prognosis.

## Discussion

EMT is a hallmark of many physiological and pathological processes contributing to morphogenesis, wound healing and cancer metastases [1]. With recently growing acknowledgement that EMT is not an ‘all-or-none’ process [1], much focus has moved to identifying ‘phenotypic stability factors’ (PSFs), i.e. players that can maintain a hybrid E/M phenotype stably, such as GRHL2, OVOL2, ΔNp63α and NUMB [6, 8–10] as well as understanding the unique traits of intermediate phenotypes [2, 27].

Here, we developed a mathematical framework to investigate the role of the transcription factor NRF2 (nuclear factor erythroid 2 related factor 2) in mediating EMT. NRF2 has been well-studied in the context of cellular redox status, where it acts as a regulator in response to diverse forms of stress [28], for instance by activating antioxidant response within the cell to respond to oxidative stress, thus promoting cell survival [29]. However, its role in mediating clustered cell migration – a proposed outcome of partial EMT [1] – has been only recently explored [12, 30].

These recent observations drove the development of our computational model which predicts that NRF2 can act as a PSF for the hybrid E/M phenotype. This result agrees well with the proposed role of NRF2 in mediating collective cell migration during wound healing [12], and its implications in restricting a complete EMT in pulmonary fibrosis [13]. A hybrid E/M or a partial EMT phenotype has been recently suggested to be potentially more aggressive than a complete EMT [1, 2]. Thus, the association of high levels of NRF2 with accelerated metastasis [31–33] and poor patient survival further [34] lends support to the idea of NRF2 acting as a PSF. Interestingly, knockdown of different PSFs independently lead to a complete EMT in H1975 NSCLC [6, 8, 25], hence indicating that these different pathways contribute to stabilising a hybrid E/M phenotype in a non-redundant way, and loss of each of them is sufficient for a complete EMT.

Our model further predicts that NRF2 follows a biphasic dynamics during EMT, i.e. its level increases upon EMT induction, it is maximally expressed in a hybrid E/M phenotype and decreases on completion of EMT. Therefore, NRF2 activation is proposed here as a hallmark of hybrid E/M phenotypes. While a direct and rigorous experimental validation of this prediction, along with testing how NRF2 regulates invasion in the more realistic setting of the extracellular matrix, is still lacking, preliminary evidence on NRF2 dynamics during injury-induced collective cell migration [12], and the association of NRF2 with enhanced drug resistance [35] argue in favour of this predicted dynamics, given the functional correlation between a hybrid E/M phenotype and acquired drug resistance [1]. Multi-drug resistance proteins have been identified as NRF2 target genes [36], and stable overexpression of NRF2 led to enhanced resistance of cancer cells to chemotherapeutic agents such as cisplatin, doxorubicin, and etoposide [37]. Our mathematical model predicts that such biphasic behaviour is conserved upon knockdown of NRF2 inhibition by KEAP1, but not upon knockdown of NRF2 inhibition by E-Cadherin. Thus, even in cases of mutations in KEAP1 and/or NRF2 that can impair NRF2-KEAP1 interactions, NRF2 is likely to be maximally expressed and represent a highly aggressive hybrid E/M phenotype. This asymmetry can perhaps offer an explanation for why KEAP1 and/or NRF2 mutations still result in poor prognosis [38, 39]. Importantly, NRF2 is not solely regulated via EMT. Several Cullin scaffold proteins support NRF2 ubiquitination. NRF2 can be targeted for degradation by the Cullin1/Rbx1 complex via recognition by the ubiquitin kinase adapted beta-trCP [40]. Further, Cullin4 regulates NRF2 independently from Keap1 via the DDB1/WDR23/Cullin4 pathway [41]. These further layers of regulation, however, do not modify the coupling between NRF2 and EMT that gives rise to the predicted biphasic behaviour of NRF2.

Our earlier efforts at quantifying the association between EMT and stemness have predicted that cells in hybrid E/M states are more inclined to gain stem-like properties [27, 42, 43], a prediction that has been recently validated *in vitro* and *in vivo* [2, 44, 45]. Reinforcing these predictions, NRF2 has been related to cancer stem-like traits and chemo-resistance [35]. Remarkably, a recent study reported a strong NRF2-mediated antioxidant response in hybrid E/M-like breast cancer stem cells (BCSC), and a significant association of NRF2 with poor prognosis [46]. These experimental observations resonate well with the worse clinical outcome and an association of a hybrid E/M phenotype with high NRF2 expression, and support the increasingly accepted notion that a hybrid E/M phenotype, rather than a completely mesenchymal one, may correlate with a higher metastatic potential [1].

Furthermore, NRF2 overexpression enhanced both epithelial and mesenchymal traits in RT4 bladder cancer cells that were already co-expressing E-cadherin and ZEB1. Previous experiments have shown the effect of NRF2 on TGF-β1-driven EMT in epithelial cells [13, 16, 47, 48], and how initial levels of NRF2 can impact the response of cells to TGF-β1-driven EMT [49]. However, our study illustrates a different role for NRF2 in modulating EMT when NRF2 is pharmacologically over-expressed in hybrid E/M cells. Therefore, the effects of NRF2 may depend on the intrinsic EMT phenotype of cells. In other words, perturbing NRF2 signalling could have different effects depending on the position of the considered cells along the EMT spectrum.

To further investigate the implications of NRF2 – or any other PSF – in promoting clustered cell migration, future modelling efforts should be devoted toward integrating mechanics of cell motility with the regulatory biochemical signalling pathways. Such efforts will offer a more reliable quantitative relationship between NRF2 signalling axis and the aggressiveness of the disease.

## Materials and Methods

### Mathematical model

The level of a protein, mRNA or micro-RNA (X) is described via a chemical rate equation that assumes the generic form:

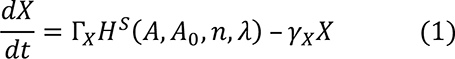

In (eq. 1), the first term on the right-hand side (Γ_*X*_*H*^*S*^(*A*, *A*_0_, *n*, *λ*)) represents the production rate of X expressed as a product of a basal production rate constant (Γ_*X*_) and a function (*H*^*S*^(*A*, *A*_0_, *n*, *λ*)) that represents transcriptional/translational/post-translational regulations emerging from interactions with other components of the system. The second term (– *γ*_*X*_*X*) is the degradation rate of X given by the product of a degradation rate constant (*γ*_*X*_) and the number of molecules (*X*) (assuming first-order kinetics).

Specifically, an activator/inhibitor (A) of X alters the production rate constant according to a shifted Hill function [18], defined as:

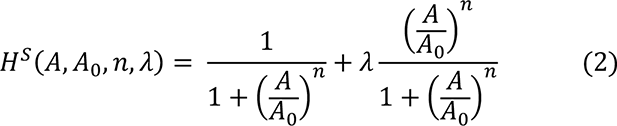

where the first argument (*A*) of the Hill function is the number of molecules of the inhibitor/activator while the second argument (*A*_0_) is the half-maximal concentration parameter. Additionally, the Hill function is characterized by the Hill coefficient *n* that relates to how steep the regulation is, and the fold change *λ* (*λ* < 1 for inhibition, *λ* > 1 for activation). If multiple components modulate X, the production rate is a product of Hill functions. The NRF2-EMT model describes the dynamics of the circuit depicted in Fig. 1A (SI section 1) by extending the EMT circuit model proposed by Lu et al [18]. The newly-introduced parameters for this model are taken from relevant references [13, 15, 16, 50–52]. A qualitative rationale for parameter estimation is provided in SI section 2.

### Analysis of clinical data

For all the examined datasets, the group of patients was divided into two sub-groups based on median NRF2 level. Kaplan-Meier analysis, i.e. hazard ratios and confidence intervals for NRF2 and its downstream targets for the two subgroups, was obtained with PROGgeneV2 software [53].

### EMT score quantification

The EMT Metric [24] assigns to each sample categorical probabilities of membership in {E, E/M, M} by performing logistic regression against a collection of EMT-relevant predictor transcripts and cross-platform normalizer transcripts. The output of sample *i* may be represented by an ordered triple of probabilities *S*_*i*_ = (*P*_*E*_, *P*_*E*/*M*_, *P*_*M*_). Categorization is assigned based on the maximal value of this ordered triple. *S*_*i*_ is then projected onto [0,2] by use of the EMT metric. The metric places epithelial and mesenchymal samples close to 0 and 2, respectively; while maximally hybrid E/M samples are assigned values close to 1.

### Cell culture and treatments

H1975 cells were cultured in RPMI 1640 medium containing 10% fetal bovine serum (FBS) and 1% penicillin/streptomycin cocktail (Thermo Fisher Scientific, Waltham, MA). Cells were transfected at a final concentration of 50 nM siRNA using Lipofectamine RNAiMAX (Thermo Fisher Scientific) according to the manufacturer’s instructions using following siRNAs: siControl (Silencer Select Negative Control No. 1, Thermo Fisher Scientific), siNRF2 #1 (n290469, Silencer Select siRNA, Invitrogen), siNRF2 #2 (n290470, Silencer Select siRNA, Invitrogen).

RT4-WT and Crispr/Cas9 NRF2-KO cells were cultured in McCoy’s 5A medium containing 10% FBS and 0.1% Gentamicin (Fisher Scientific, Hampton, NH). Cells were maintained at 37 °C in 5% CO_2_ and media was refreshed every 2 days. Crispr-Cas9 knockout cell pool RT4-NRF2-KO was obtained from Synthego, CA. The following gRNA targeting exon 2 was used for NFE2L2-KO: AUUUGAUUGACAUACUUUGG. Knockout cells show a predicted functional knockout of 63% which was confirmed by Synthego through RT-qPCR showing 75% editing efficiency post expansion at passage 4 (data not shown). All experiments were done between passages 5-8 for the Crispr/Cas9 NRF2-KO cells. All migration assays and immunocytochemistry experiments were performed in polystyrene 24-well plates.

### RT-PCR

Total RNA was isolated following manufacturer’s instructions using RNAeasy kit (Qiagen). cDNA was prepared using iScript gDNA clear cDNA synthesis kit (Bio-Rad). A TaqMan PCR assay was performed with a 7500 Fast Real-Time PCR System using TaqMan PCR master mix, commercially available primers, and FAM™-labeled probes for CDH1, VIM, ZEB1, NRF2, SNAIL, and VIC™-labeled probes for 18S, according to the manufacturer’s instructions (Life Technologies). Each sample was run in biological and technical triplicates. Ct values for each gene were calculated and normalized to Ct values for 18S (ΔCt). The ΔΔCt values were then calculated by normalization to the ΔCt value for control.

### Western Blotting Analysis and Immunofluorescence

H1975 cells were lysed in RIPA lysis assay buffer (Pierce) supplemented with protease and phosphatase inhibitor. The samples were separated on a 4–15% SDS-polyacrylamide gel (Biorad). After transfer to PVDF membrane, probing was carried out with primary antibodies and subsequent secondary antibodies. Primary antibodies were purchased from the following commercial sources: anti-CDH1 (1:1000; Cell Signaling Technology), anti-vimentin (1:1000; Cell Signaling Technology), anti-Zeb1 (1:1000; Cell Signaling Technology), anti-SNAIL (1:1000; Cell Signaling Technology), anti-NRF2 (1:1000; Abcam), and anti-GAPDH (1:10,000; Abcam). Membranes were exposed using the ECL method (GE Healthcare) according to the manufacturer’s instructions. The band intensity of three independent experiments were quantified using Image-J software and then normalized against the loading control. For immunofluorescence, cells were fixed in 4% paraformaldehyde, permeabilized in 0.2% Triton X-100, and then stained with anti-CDH1 (1:100; Abcam) and anti-vimentin (1:100; Cell Signaling Technology). The primary antibodies were then detected with Alexa conjugated secondary antibodies (Life technologies). Nuclei were visualized by co-staining with DAPI.

### Immunocytochemistry

RT4-WT and Crispr/Cas9 NRF2-KO cells were washed with warmed PBS twice, followed by fixation with chilled 4% paraformaldehyde (Sigma) in PBS for 15 min, all reagents were kept cold past this point and incubation was performed at room temperature unless stated otherwise. Permeabilization was performed with 1% Triton X-100 in PBS for 10 minutes followed by a blocking step with 3% bovine serum albumin (BSA) in PBS for 30 minutes. The cells were incubated overnight at 4 °C with the primary antibodies and then incubated in the dark for 2 hours against the secondary antibodies. Primary antibodies used were NRF2 (1:50; cat. # sc-13032, Santa Cruz Biotechnology), HO-1 (1:100; cat. # ab13248, Abcam), E-cadherin (1:50; cat. # M3612, Dako), and ZEB1 (1:50; cat # ab124512, Abcam). Secondary antibodies used were Alexa conjugated secondary antibodies (1:250, Life technologies). The antibodies were all diluted in 3% BSA solution. Wells were washed 3 times with PBS in between each step. Cells were examined using a laser scanning confocal microscope (Leica TCS SP8; Leica Microsystems, Wetzlar, Germany).

### Wound-Healing assay

Scratch wound-healing assay was performed to determine cell migration using confluent cultures (80%−90% confluence). Briefly, cells (1 × 105 cells/ml) were seeded in 6-well tissue culture plate and grown confluent. Cells were starved for 24 hours using 0.2% serum in growth media to minimize cell proliferation. Next day, the confluent monolayer was scratched with a sterile p200 pipet tip and media replaced by complete growth media. Images were acquired at 0 and 16 hours; the experiments were repeated 3 times. Images of the scratch wounds were taken and measured by Image-J software to calculate the mean and standard deviation. Each group was compared with the control group. Cell migration was expressed as the migration rate: (original scratch width - new scratch width)/original scratch width ×100%.

In RT4-WT and Crispr/Cas9 NRF2-KO cells, scratch wound-healing assay was performed to determine cell migration using confluent cultures. Briefly, at 100% confluency, the monolayer was scratched with a sterile 1 mL pipet tip and media replaced by complete growth media. Cells were washed with warm 1x PBS before and after wounding. Images were acquired at 0, 24 and 48 hours; the experiments were repeated at least 3 times and Image-J software was used to calculate the mean and standard deviation. Cell migration was expressed as the migration rate in microns per hour (μm/h): (original scratch width – final scratch width)/time.

### Trans-well migration assay

H1975 cells were grown in 6-well plates and treated with siRNAs for NRF2 for 24 hours. After 48hrs of siRNA knockdown, cell monolayers were harvested, counted and cell concentration was adjusted to 2 × 10^4^ viable cells/200 μl of serum free medium. The cell suspension was seeded on top of a Transwell insert with 0.8 μm pore diameter (Millipore) and placed on a 24 well cell culture plate. At the bottom of the Transwell insert, 10% fetal bovine serum was added as chemo-attractant and plate was incubated at 37°C for 18 hours. Medium was aspirated from assay plate and Transwell inserts. Cells that did not migrate were removed by gently swabbing the inside of each insert using cotton swabs. After 3 washes with PBS, cells were fixed and stained with a 0.5% crystal violet solution for 10 minutes. After the cells have been stained, the inserts were washed thoroughly with water until the water runs clear. Inserts were dried completely before visualizing with a microscope. Cell numbers were counted using an inverted microscope at ×200 magnification. The experiment was repeated 3 times and statistically analysed with 5 fields of view, and the mean values were taken as the migratory cell number.

## Author’s contribution

FB and MKJ conceived the research; FB (mathematical model, RACIPE simulation), SCT (H1975 NSCLC) and SAVM (RT4) developed the methodology; FB, JTG and MKJ analysed clinical datasets; all authors discussed results and participated in the preparation of the manuscript.

## Acknowledgements

Federico Bocci thanks Dongya Jia and Bin Huang for helpful conversations about RACIPE simulations.

