## Supplementray information for "NRF2 activates a partial Epithelial-Mesenchymal Transition and is maximally present in a hybrid Epithelial/Mesenchymal phenotype"

**Running title:** NRF2 is a hallmark of a hybrid E/M phenotype

Federico Bocci*^,1,2^, Satyendra C Tripathi*^,6^, Samuel A Vilchez Mercedes*^,8^, Jason T George^1,3,7^, Julian P Casabar^6^, Pak K Wong^8,9^, Samir M Hanash^6^, Herbert Levine^1,2,3,4,#^, José N Onuchic^1,2,4,5#^, Mohit Kumar Jolly^1,10,#^

^1^Center for Theoretical Biological Physics, Department of ^2^Chemistry, ^3^Bioengineering, ^4^Physics and Astronomy, ^5^Biosciences, Rice University, Houston, TX, USA;

^6^Department of Clinical Cancer Prevention, UT MD Anderson Cancer Center, Houston, TX, USA; ^7^Medical Scientist Training Program, Baylor College of Medicine, Houston, TX, USA;

^8^Department of Biomedical Engineering, The Pennsylvania State University, University Park, PA, USA; ^9^Department of Mechanical Engineering and Department of Surgery, The Pennsylvania State University, University Park, PA, USA; ^10^ Current address: Centre for BioSystems Science and Engineering, Indian Institute of Science, Bangalore, India

*These authors equally contributed to this work.

^#^corresponding authors:

Herbert Levine:

José N Onuchic:

Mohit Kumar Jolly:

1. **NRF2-EMT model**

The computational framework for NRF2-EMT circuit describes the dynamics of the molecular species of the EMT regulatory circuit (miR-34, miR-200, Snail, Zeb), NRF2 and the molecular players that participate in the connection between NRF2 and the EMT circuit (KEAP1, E-cadherin) as shown in Fig. 1A. This framework extends the mathematical model of EMT circuit developed earlier (1). The temporal dynamics of the species of the circuit (miR-34: W, miR-200: Y, Snail: S, Zeb: Z, KEAP1: K, E-cadherin: E, NRF2: X) are modeled via a system of ordinary differential equations:

$$\frac{d\mu_{34}}{dt}=g_{\mu_{34}}H^{S}\left( S \right)H^{S}\left( Z \right)-g_{S}H^{S}\left( S \right)H^{S}\left( I \right)H^{S}\left( I_{ext} \right)P_{y}\left( \mu_{34},2 \right) -\gamma_{\mu_{34}}\mu_{34}$$

$$\frac{d\mu_{200}}{dt}=g_{\mu_{200}}H^{S}\left( Z \right)H^{S}\left( S \right)-g_{Z}H^{S}\left( Z \right)H^{S}\left( S \right)P_{y}\left( \mu_{200},6 \right) -\gamma_{\mu_{200}}\mu_{200}$$

$$\frac{dZ}{dt}={k_{P}g}_{Z}H^{S}\left( Z \right)H^{S}\left( S \right)P_{l}\left( \mu_{200},6 \right) -\gamma_{Z}Z$$

$$\frac{dS}{dt}={k_{P}g}_{S}H^{S}\left( X \right)H^{S}\left( S \right)H^{S}\left( I_{ext} \right)P_{l}\left( \mu_{34},2 \right) -\gamma_{S}S$$

$$\frac{dK}{dt}={k_{K}H}^{S}\left( \mu_{200} \right) -\gamma_{K}K$$

$$\frac{dE}{dt}={k_{E}H}^{S}\left( Z \right) -\gamma_{E}E$$

$$\frac{dX}{dt}={k_{X}H}^{S}\left( K \right)H^{S}\left( E \right) -\gamma_{X}X$$

Here, $g_{\mu_{200}}$, $g_{\mu_{34}}$, $g_{Z}$and $g_{S}$ are the basal transcription rates of miR-200, miR-34, Zeb and Snail, while $k_{P}$ is the translation rate for Zeb and Snail. The production of KEAP1, E-cadherin and NRF2 is modeled via a single production rate that includes both transcription and translation ($k_{K},k_{E}, k_{X}$, respectively). Any species $i$ has its own degradation rate $\gamma_{i}$. Transcriptional/translational activation/inhibition in the circuit are modeled via shifted Hill functions:

$$H^{S}\left( A,A_{0},n,\lambda\right)= \frac{1}{1+\left( \frac{A}{A_{0}} \right)^{n}}+\lambda\frac{\left( \frac{A}{A_{0}} \right)^{n}}{1+\left( \frac{A}{A_{0}} \right)^{n}}$$

where the first argument ($A$) of the Hill function is the molecular player that poses the regulation while the second argument ($A_{0}$) is the half-maximal concentration parameter in a Hill function. Additionally, the Hill function is characterized by the Hill coefficient $n$ that relates to how steep the regulation is with respect to the concentration of activator/inhibitor and the fold change $\lambda$ ($\lambda<1$ for inhibition, $\lambda>1$ for activation). For simplicity, only the first argument of the function $H^{S}$ has been displayed in the equations.

The functions $P_{l}(\mu,n)$ represent the post-translational inhibition that a microRNA species $\mu$ exerts on its target, where $n$ is the number of corresponding available binding sites on the target mRNA. $P_{y}(\mu,n)$ describes the decrease in the level of microRNA due to the degradation of the microRNA/mRNA complex. These functions are rigorously derived in the Supplementary Information of Lu et al. (1). The production/degradation rates of NRF2, KEAP1 and E-cadherin and the threshold for activation/inhibition, Hill coefficient and fold-change for all NRF2-related interactions are presented in Table 1. All parameters of the pre-existing model of the EMT core network were not modified, and could be found in Lu et al.(1). Also, the parameters related to the mutual inhibition between Zeb and E-cadherin were taken from Mooney et al(2).

2. **Parameters estimation**

Production and degradation rates of NRF2, E-cadherin and KEAP1 were chosen to achieve a typical cellular baseline (i.e. in absence of any activation or inhibition) of hundreds of thousands of molecules(3) in the absence of any activation/inhibition (specifically, we chose a level of $500000 molecules$), the typical number of protein molecules that can be expected in a cell line. Under such conditions, the cellular level of any species $i$ is simply given by $g_{i}/\gamma_{i}$. The degradation rate was taken as $\gamma_{i}=0.1 h^{-1}$ based on a half-life of around 7 hours(4), so the corresponding production rate (which considers both transcription and translation) is $g_{i}=50000 molecules/h$. The fold change $\lambda$ in the production rate of any species was inferred from literature (references are indicated in Table 1). All the newly introduced Hill coefficients were assumed to be 2. The half-maximal concentration (also known as Hill function threshold) for every Hill function was chosen to enable a tangible effect of the regulator, given the baseline levels of the various species.

**Supplementary Tables**

| Group | Parameter | Value | Reference |
| --- | --- | --- | --- |
| G1: NRF2 basal production/degradation | $g_{Nrf2}$ | $5{10}^{4}molec/h$ | Zeiler et al(3) |
|  | $\gamma_{Nrf2}$ | $0.1 h^{-1}$ | Eden et at(4) |
| G2: KEAP1 basal production/degradation | $g_{K}$ | $5{10}^{4}molec/h$ | Zeiler et al(3) |
|  | $\gamma_{K}$ | $0.1 h^{-1}$ | Eden et at(4) |
| G3: E-cadherin basal production/degradation | $g_{E}$ | $5{10}^{4}molec/h$ | Mooney et al(2) |
|  | $\gamma_{E}$ | $0.1 h^{-1}$ | Mooney et al(2) |
| G4: inhibition of Snail by NRF2 | $X_{0S}$ | $1.0{10}^{6}molec$ | estimated |
|  | $\lambda_{X,S}$ | $0.67$ | Zhou et al(5) |
| G5: inhibition of NRF2 by E-cadherin | $E_{0X}$ | $2.5{10}^{5}molec$ | estimated |
|  | $\lambda_{E,S}$ | $0.33$ | Kim et al(6) |
| G6: inhibition of KEAP1 by miR-200 | $W_{0K}$ | $5.0{10}^{3}molec$ | estimated |
|  | $\lambda_{W,K}$ | $0.1$ | Eades et al(7) |
| G7: inhibition of NRF2 by KEAP1 | $K_{0X}$ | $2.5{10}^{5}molec$ | estimated |
|  | $\lambda_{K,X}$ | $0.33$ | Eades et al(7) |

Supplementary Table S1. Parameters of the NRF2-EMT model. First column classifies parameters by groups based on their functional role in the circuit.

**Supplementary figures**

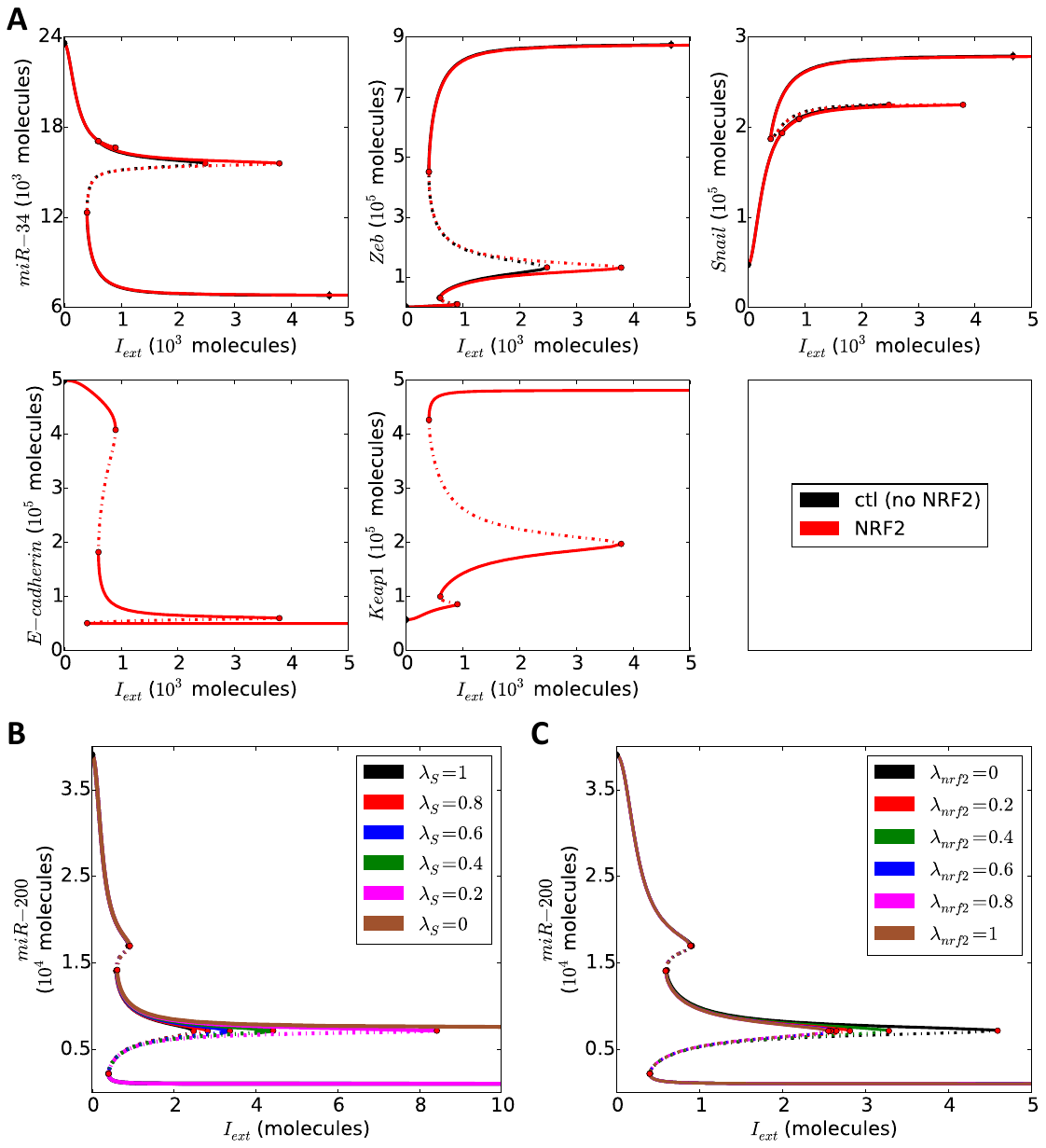

Figure S1. **(A)** Bifurcation diagram of miR-34, ZEB, SNAIL, E-Cadherin and KEAP1 as a function of external EMT-inducing signal ($I_{ext}$). Control case (black curve) and NRF2-dependent case (red curve) are very similar in miR-34, ZEB and SNAIL with the exception that the range of stability of hybrid E/M state (i.e. the range of $I_{ext}$ values for which the hybrid E/M state can be observed) is larger in the NRF2-dependent case. E-Cadherin and KEAP1 are computed only in the NRF2-dependent case because they are not included in the control EMT model. **(B)** Bifurcation diagram of miR-200 for different fold-change values ($\lambda_{S}$) of SNAIL production rate constant due to NRF2 inhibition. **(C)** Bifurcation diagram of miR-200 for different fold-change values ($\lambda_{nrf2}$) of NRF2 production rate constant due to the inhibition by E-cadherin and KEAP1. **(B-C)**: increasing the inhibition (i. e. decreasing the $\lambda$s from 1 to 0) increases the stability range of the hybrid E/M state but does not alter the stability of the E and M states.

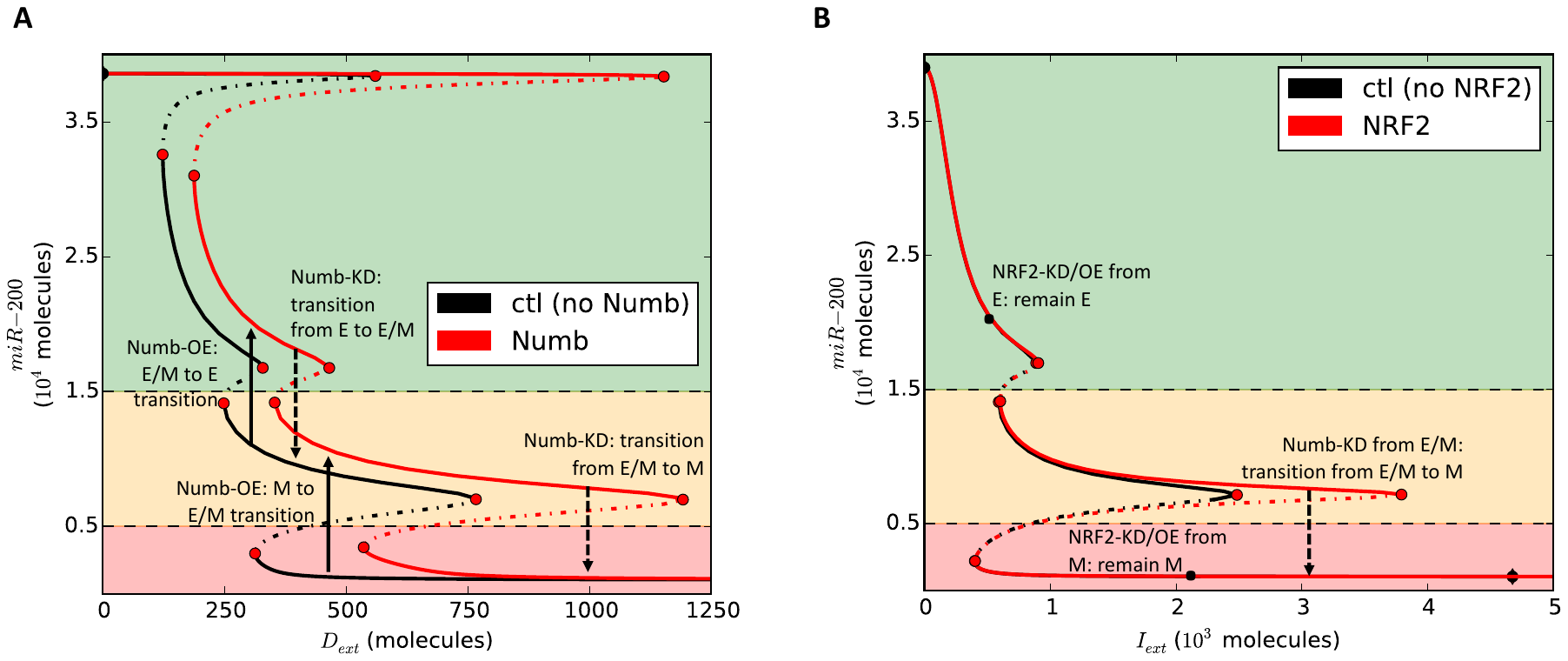

Figure S2. **(A)** Bifurcation diagram of miR-34 as a function of external ligand Delta ($D_{ext}$) in the model of EMT-Notch-Numb signalling adapted from Ref. 8. Knockdown of Numb from an Epithelial or hybrid E/M phenotype can result in a transition to hybrid E/M or Mesenchymal respectively. Conversely, expression of Numb from hybrid E/M or M can result in a transition to E or hybrid E/M, respectively. **(B)** Bifurcation diagram of miR-34 as a function of external EMT inducer $I_{ext}$ (same as figure 1A). Knockdown of NRF2 from a hybrid E/M phenotype results in a transition to a M phenotype, but NRF2-KD from an E or M phenotype has no predicted effect in the model.

**E-cadherin**

**ZEB1**

**T= 1 day**

**T= 16 days**

**T=28 days**

**A**

**B**

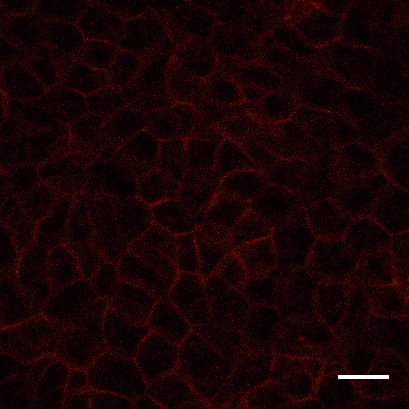

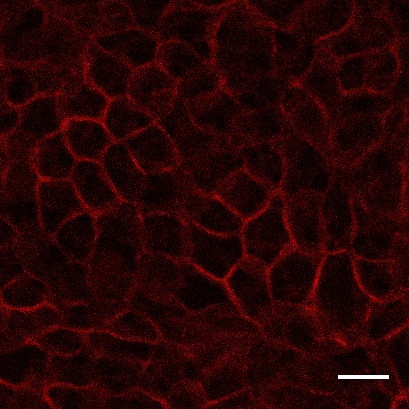

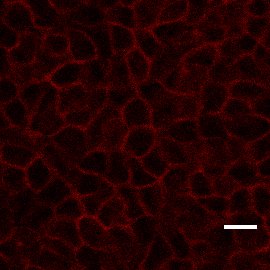

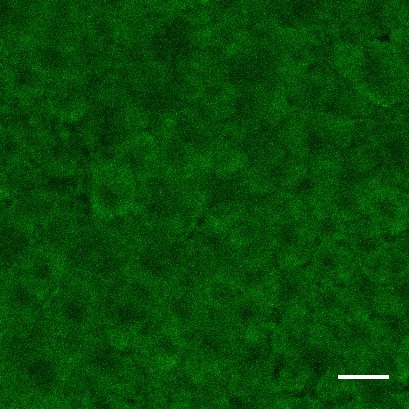

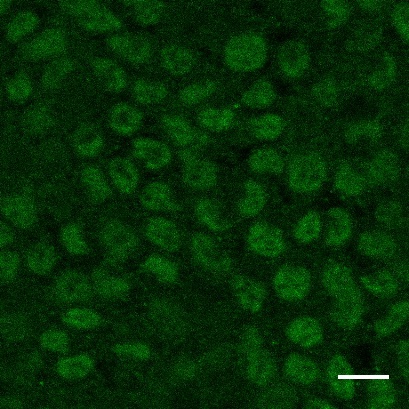

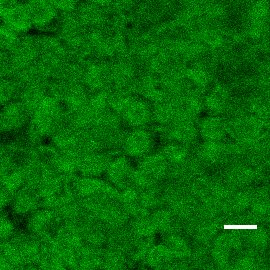

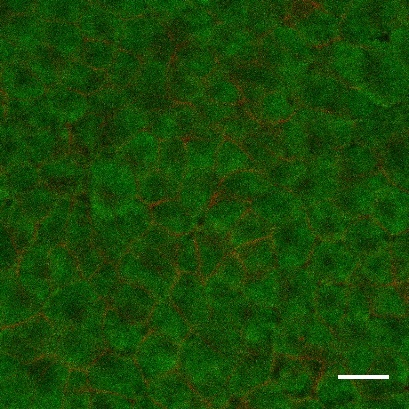

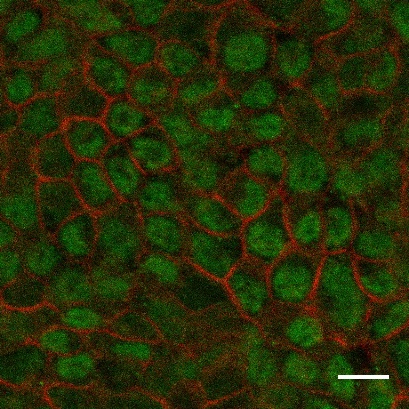

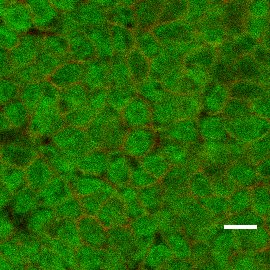

**Merged**

**C**

Figure S3. RT4 cells demonstrate a stable hybrid E/M phenotype over multiple passages (T denotes number of days between experiments, and cells were passaged every 4-6 days). Immunocytochemical staining of epithelial marker E-cadherin **(A)**, mesenchymal marker ZEB1 **(B)**, and a merged view **(C)** indicating stable co-expression of both markers over multiple passages. All experiments were performed between passages 5 and 20. Scale bar: 20 μm.

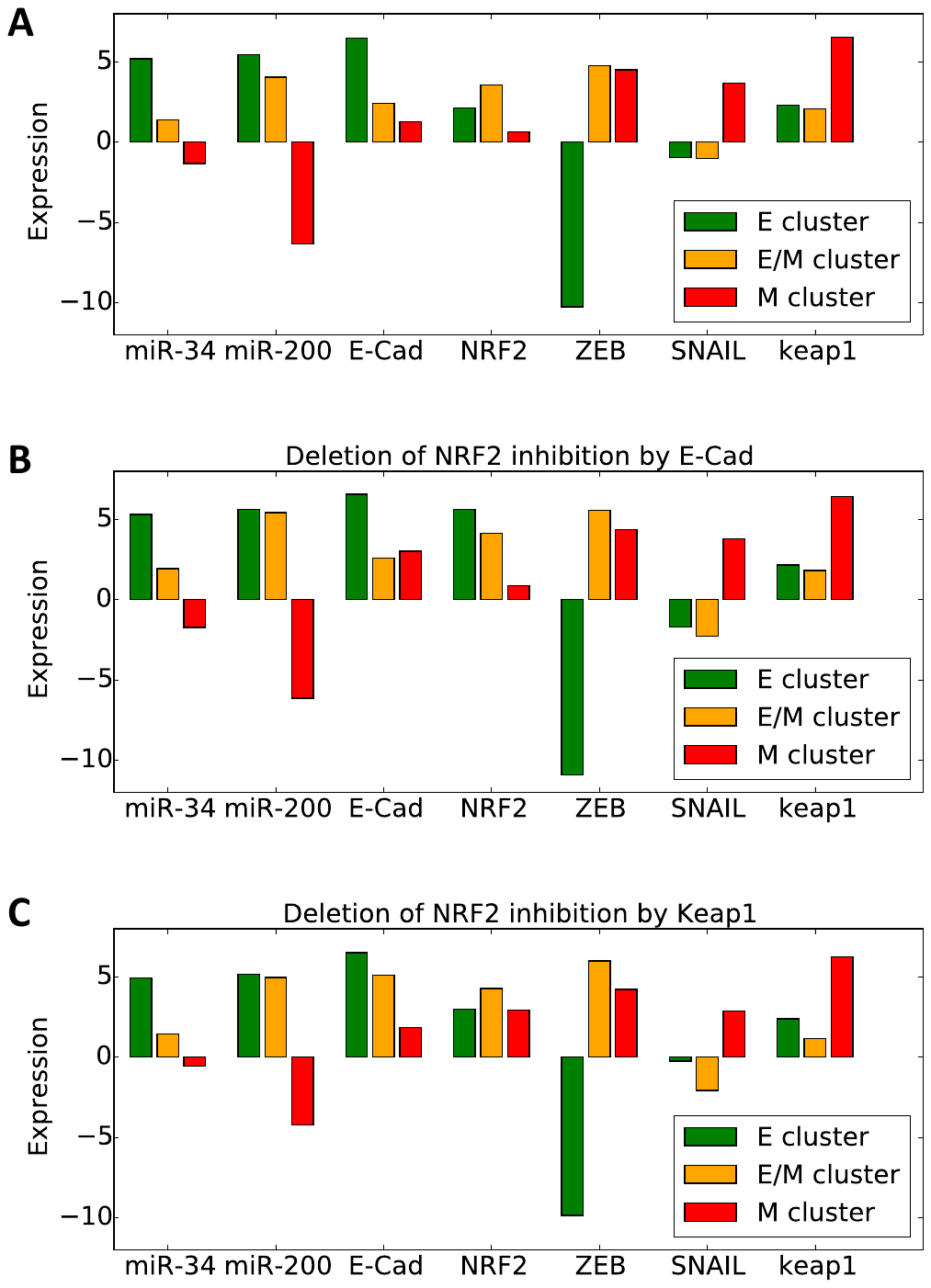

Figure S4. Average expression level of all the model’s variables in the three groups obtained from the clustering 10000 RACIPE simulations of the EMT-NRF2 circuit for **(A)** All considered interactions (same as in Fig. 3); **(B)** all interactions but the inhibition of NRF2 by E-Cadherin and; **(C)** all interactions but the inhibition of NRF2 by KEAP1. NRF2 maintains the characteristic high level in the hybrid E/M cluster when KEAP1-NRF2 interaction is knocked-down. Conversely, NRF2 levels become directly correlated to miR-200 levels when the NRF2-E-Cadherin interaction is knocked down.

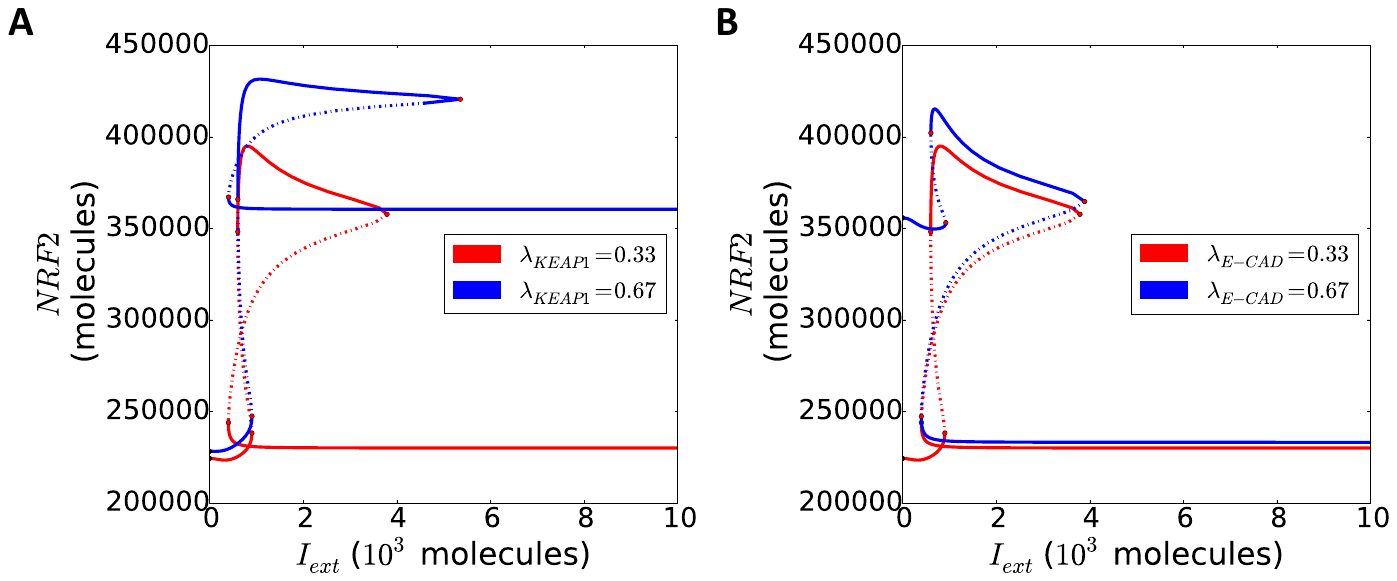

Figure S5. **(A)** Bifurcation diagram of NRF2 levels as a function of EMT-inducing signal ($I_{ext}$) for the original fold-change value in KEAP1-mediated inhibition of NRF2 (red) and for a higher (i.e. weaker inhibition) value (blue). **(B)** Same as in A, but varying E-Cadherin-mediated inhibition of NRF2 instead.

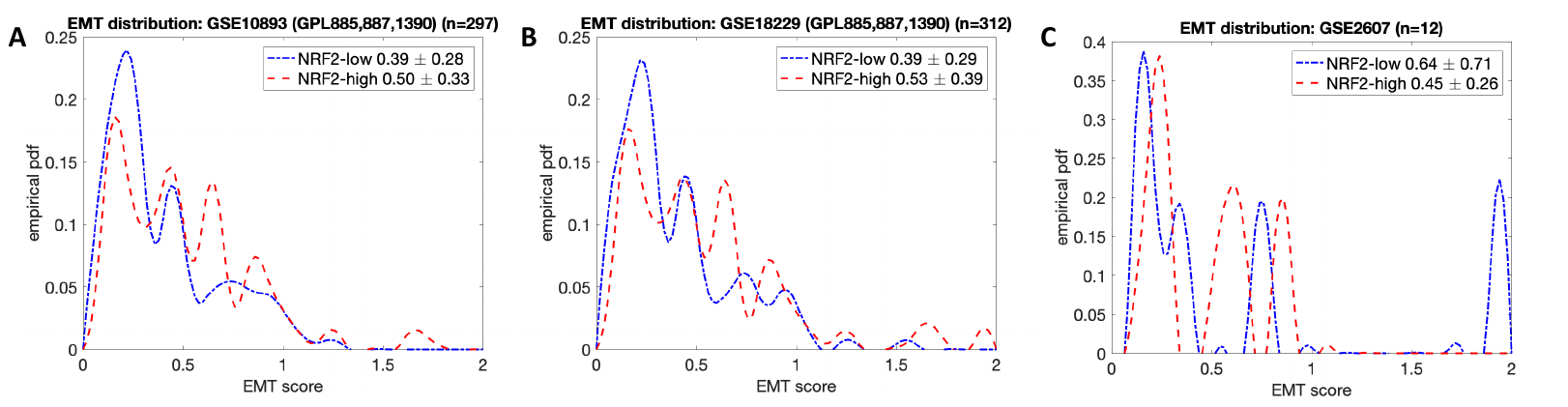

Figure S6. EMT score distribution for the patient from the clinical datasets GSE10893 **(A)**, GSE18229 **(B)** and GSE2607 **(C)**. Blue and red curves correspond to ‘low NRF2’ group and ‘high NRF2’ group, respectively. Panels indicate the average EMT score and standard deviation within each group. The probability of a hybrid E/M EMT score corresponds to the area underlying the distribution in the interval [0.5, 1.5]. This area is larger for the ‘high NRF2’ red curves, as quantified in Fig. 4E. Therefore, a patient is more likely to have a hybrid E/M EMT score if he/she belongs to the ‘high NRF2’ group as compared to the ‘low NRF2’ group.
